# Intraspecific variation at the free-living stage of a soil-borne pathogen is organized into an exploration syndrome associated with infectivity

**DOI:** 10.64898/2026.10.09.757656

**Authors:** Margot Trinquier, Marie-Laure Pilet-Nayel, Clément Lavaud, Baptiste Mayjonade, Sébastien Carrère, Jérôme Gouzy, Hervé Philippe, Delphine Legrand, Elodie Gaulin

## Abstract

Many pathogens spend a substantial portion of their life cycle outside the host tissues. During its free-living stage, the root legume-plant pathogen *Aphanomyces euteiches* persists in soil either as non-motile resting oospores or actively explores the environment through mycelium and motile zoospores. These two latter stages are critical for survival, dispersal and host encounter. However, the extent of intraspecific variation during these phases remains poorly explored, and it is still unknown whether they are organized into phenotypic syndromes contributing to infection success. Here, we measured eight free-living phenotypic traits associated with mycelial and zoospore stages across 23 *A. euteiches* strains from distinct geographic regions (USA, Europe) whose genomes were sequenced. We tested whether traits are organized into a phenotypic syndrome linked to either geographic origin, genomic diversity or infectivity. The microbial strains exhibited substantial intraspecific phenotypic variation that we proposed as an ‘exploration syndrome’ linked to host search. Beyond this general syndrome, we detected sub-structure according to strains’ infectivity, geographic origin and genetic group. This suggests that the free-living stage of pathogens may contribute to disease potential. Our results highlight the importance of integrating functional trait approaches into plant pathology and suggest that the identified ‘exploration syndrome’ may serve as a predictor of infectivity in soilborne oomycete pathogens.

## INTRODUCTION

Plant-pathogen interactions play a pivotal role in ecosystems functioning and in agroecosystem productivity. Plant pathogens are responsible for annual crop losses of 20–40% and economic damages exceeding US$220 billion worldwide (FAO 2019; Savary *et al*. 2019). These impact highlights the urgent need to better understand the earliest stages of infection, a process influenced by pathogen life-history strategies. Among plant pathogens, soil-borne species are challenging because they spend a substantial portion of their lifecycle outside host tissues; persisting, dispersing and encountering hosts during a free-living stage (Caraco & Wang 2008). This stage is critical for pathogens survival and infection success, yet its role in shaping pathogenicity remains largely unexplored.

Agroecosystems, characterized by high-density monocultures grown continuously over large spatial scale, provide an ideal environment for the emergence and evolution of pathogens by facilitating host specialization, rapid adaptation and the expansion of pathogen populations (McDonald & Stukenbrock 2016). These processes are further accelerated by human activities that enable global dissemination of pathogens. Among the most destructive pathogens are oomycetes, filamentous eukaryotic microorganisms belonging to the kingdom *Stramenopila.* These organisms occupy diverse terrestrial and aquatic habitats and include several of the world’s most devastating plant pathogens, such as *Phytophthora infestans*, the causal agent of potato late blight (Kasteel *et al*. 2023), and *Aphanomyces euteiches*, responsible for root rot in legumes (Beakes *et al*. 2012; Gaulin *et al*. 2007; Wu *et al*. 2018).

Among root-infecting oomycetes, *A. euteiches* is one of the most destructive pathogen of grain legumes worldwide. Despite substantial progress in the identification of resistant pea genotypes and in deciphering the molecular basis of pathogenicity (Camborde *et al*. 2019, 2022; Gaulin *et al*. 2018; Leprévost *et al*. 2023; Leprévost & Pilet-Nayel 2026), our understanding of the pathogen remains largely based on fine-scale root interactions and a limited number of reference strains, leaving the extent and ecological significance of its intraspecific diversity largely unexplored.

First described in North America in the 1920s (Jones *et al*. 1925), *A. euteiches* has since been recorded in pea-growing regions across Europe (notably in France and Norway), Canada, North America, Australia, New Zealand and Japan (Labrousse 1933; Sundheim 1972). In France, the species emerged as a significant economic threat in the 1990s (Wicker & Rouxel 2001) and remains one of the most damaging pathogens in legume production systems. Considerable efforts have been devoted to identify resistance QTLs in pea and to deciphering the molecular mechanisms underlying plant–pathogen interactions (Desgroux *et al*. 2016; Lavaud *et al*. 2025; Leprévost *et al*. 2025; Pilet-Nayel *et al*. 2005) and the molecular determinants of pathogenicity (Camborde *et al*. 2019, 2022; Gaulin *et al*. 2018). Nevertheless, as for many plant-pathogen systems, research on *A. euteiches* has largely relied on a limited number of reference strains, such as RB84 and ATCC201684 (France, pathotype I) and Ae109 (USA, pathotype III), which have served as the primary models for genomic and pathogenicity studies. These strains were primarily selected because they exhibit distinct pathogenicity profiles on legume hosts and represent contrasting geographic origins (Kiselev *et al*. 2022; Leprévost *et al*. 2025; Moussart *et al*. 2024; Quillévéré-Hamard *et al*. 2018). However, if the first studies of pathogenicity profiles indeed showed that French strains were mostly classified as pathotype I (Quillévéré-Hamard *et al*. 2018), North American strains displayed greater diversity, including pathotypes I, III, V, VI, and X (Hamon *et al*. 2011; Malvick & Percich 1998; Sivachandra Kumar *et al*. 2021; Wicker & Rouxel 2001). This suggests that intraspecific variability could be a key neglected aspect in the study of *A. euteiches*. Accordingly, recent studies have revealed a much broader genetic and pathogenic diversity than generally thought, including geographically structured populations in the USA and France (Le May *et al*. 2018), three distinct genetic groups in Europe (Kälin *et al*. 2022) with the identification of an Italian group as a potential cryptic species (Kälin *et al*. 2026) and the discovery of pathotype II in Canadian and US fields (Sivachandra Kumar *et al*. 2021). These findings strongly challenge the representativeness of model strains and emphasize the need for a more comprehensive analysis of the role of *A. euteiches* intraspecific variation in its biology and pathogenicity.

Intraspecific variation, *i.e.* the diversity within a species, is recognized as a powerful driver of population resilience, adaptation and persistence, shaping life-history strategies, dispersal dynamics and responses to environmental changes at population, community and ecosystem levels (Bolnick *et al*. 2011; Fargeot *et al*. 2025; Gamelon *et al*. 2025; Raffard *et al*. 2019; Violle *et al*. 2012). Concurrently, trait-based approaches provide a framework for understanding the ecological roles of organisms by focusing on measurable functional traits, including morphological, physiological, behavioural and fitness-related traits, which can influence organismal performance and fitness (Violle *et al*. 2007). This framework is now widely applied in plants (Albert *et al*. 2011) and fungi (Aguilar-Trigueros *et al*. 2014; Krah & Bässler 2021) and has been recently proposed to better understand unicellular eukaryotes diversity (Jamy *et al*. 2026). This phenotypic variation can manifest as phenotypic syndromes, *i.e.*, suites of correlated traits, accounting for individual strategies and fitness (Agrawal 2020; Sinnott-Armstrong *et al*. 2022). For instance, flowers can associate particular coloration, scent or shapes to attract pollinators (pollination syndromes) (Fenster *et al*. 2004) and dispersal is generally performed by larger and more active individuals harbouring a marked locomotion-oriented morphology (dispersal syndromes) (Cote *et al*. 2017).

While trait-based approaches and the detection of phenotypic syndromes linked to pathogenicity are increasingly applied to different organisms, their application to filamentous fungi and oomycetes pathogens remain limited. Among the few existing examples, it has been shown that isolates of *Metarhizium acridum*, an entomopathogenic fungus, harbour high variability in virulence, metabolic capability and growth indicating ecological specialisation (Parker *et al*. 2023). In *Batrachochytrium dendrobatidis*, an amphibian-killing fungus, zoospore phenotypic variation (size, production, zoosporangium size) reflects local adaptation, with morphological traits influencing pathogen transmission (Lambertini *et al*. 2016). In contrast, phenotypic diversity measured in 14 strains of *Aphanomyces astaci*, a freshwater crayfish pathogen, does not correlate with virulence despite high variability in hyphal growth and sporulation rate (Francesconi *et al*. 2024).

In *A. euteiches*, the free-living stage comprises non-motile oospores, and two motile free-living propagules: filamentous hyphae and asexual biflagellate zoospores. These motile propagules ensure survival and active exploration of the soil to locate potential infection sites (Gaulin *et al*. 2007). Yet, the extent of phenotypic diversity expressed during the free-living stage of *A. euteiches* and whether it is organized into a phenotypic syndrome potentially associated with pathogenicity remain unknown.

In this study, we investigated whether a phenotypic syndrome structures the motile free-living traits of *A. euteiches*, and whether trait variation reflects its pathogenicity, geographic origin and genomic diversity. We characterised eight traits related to mycelial development, zoospore morphology and behaviour, and fitness across 23 *Ae* strains from Europe and the USA, together with their genomic diversity and their pathogenicity. The latter was estimated with two complementary methods, one adapted from the historical method defining pathotypes (Moussart et al. 2001), and a another allowing real-time monitoring of root symptoms. We tested the three following hypotheses:

1. Traits of the motile free-living stages in *A. euteiches* present substantial variation organized into a phenotypic syndrome accounting for the exploration capacity of the pathogen;
2. Beyond this general syndrome, trait variation is sub-structured according to strains’ genomic diversity and/or geographic origin;
3. Strain pathogenicity can be predicted by at least some traits expressed at the free-living stage.

By integrating phenotypic, genomic and pathogenicity data, this work aims to provide new insights into the role of free-living stage trait variation in shaping *A. euteiches* adaptive potential and disease dynamics, while providing a framework for applying trait-based approaches in plant pathology.

## MATERIALS & METHODS

### Aphanomyces euteiches collection

We used 25 strains isolated between 1984 and 2020 from infected European pea fields (France, Denmark, Sweden and Finland) and in the USA. Details on isolation and referenced paper are given in Supplementary Table 1. Notice that due to technical reasons, genomic data were not obtained for two strains and phenotypic data not generated for two other strains (see next sections). Pathotype classification of each strain was primarily extracted from published data (Sup Tab 1). Pathotype refers to distinct groups of isolates that differ in their virulence and host interactions on pea (*Pisum sativum*). In total, 11 pathotypes have been described among pea-infecting *A. euteiches* isolates, using several pathogen’s strains origin (Europe, North America, New-Zealand) and a set of pea genotypes composed of partially resistant and susceptible lines (Wicker *et al*. 2003; Wicker & Rouxel 2001). Pathotype I is the most widespread and dominant in Europe, it is highly aggressive on most pea lines, including the susceptible reference MN313, while Pathotype III is less common and was initially identified among North American isolates. It shows lower aggressiveness toward the pea line MN313, which is susceptible to pathotype I but partially resistant to pathotype III.

Strains for which we could obtain information are classified as either pathotype I or III except Di6 and Di7, which are classified as an intermediate pathotype (IN). They have been shown to exhibit virulence traits between pathotypes I and III (Moussart *et al*. 2024). For strains Fi46, 22-2 and SE37, pathotype was not determined (Sup Tab 1).

#### Strain conservation, mycelium and zoospore production

To generate a stock of biological materials from which we initiated all experiments, we used standardized *A. euteiches* culture procedures. A stock of mycelial ‘mother’ plates was grown for seven days on Corn Meal Agar (CMA, 1.7%) in petri-dishes plates (Ø 9 cm) at 22°C in the dark. Stock cultures were subsequently maintained at 10°C in the dark and used as the source for all experiments. For mycelium production, we directly used a plug (0.5mm) of a ‘mother’ plate while for zoospore production, we used a 7 days-old mycelium produced from a plug of a ‘mother’ plate.

Sporulation of zoospores were obtained from 20 plugs (0.5mm) incubated in 20 ml of PG medium (2% bactopeptone, 0.4% glucose) at 22°C during 48h in the dark (Fig 1A). The plugs were then washed four times in sterile Volvic® water (30ml) every two hours, before a final incubation (overnight at 22°C in dark) in sterile Volvic® water (10ml).

**Figure 1.**
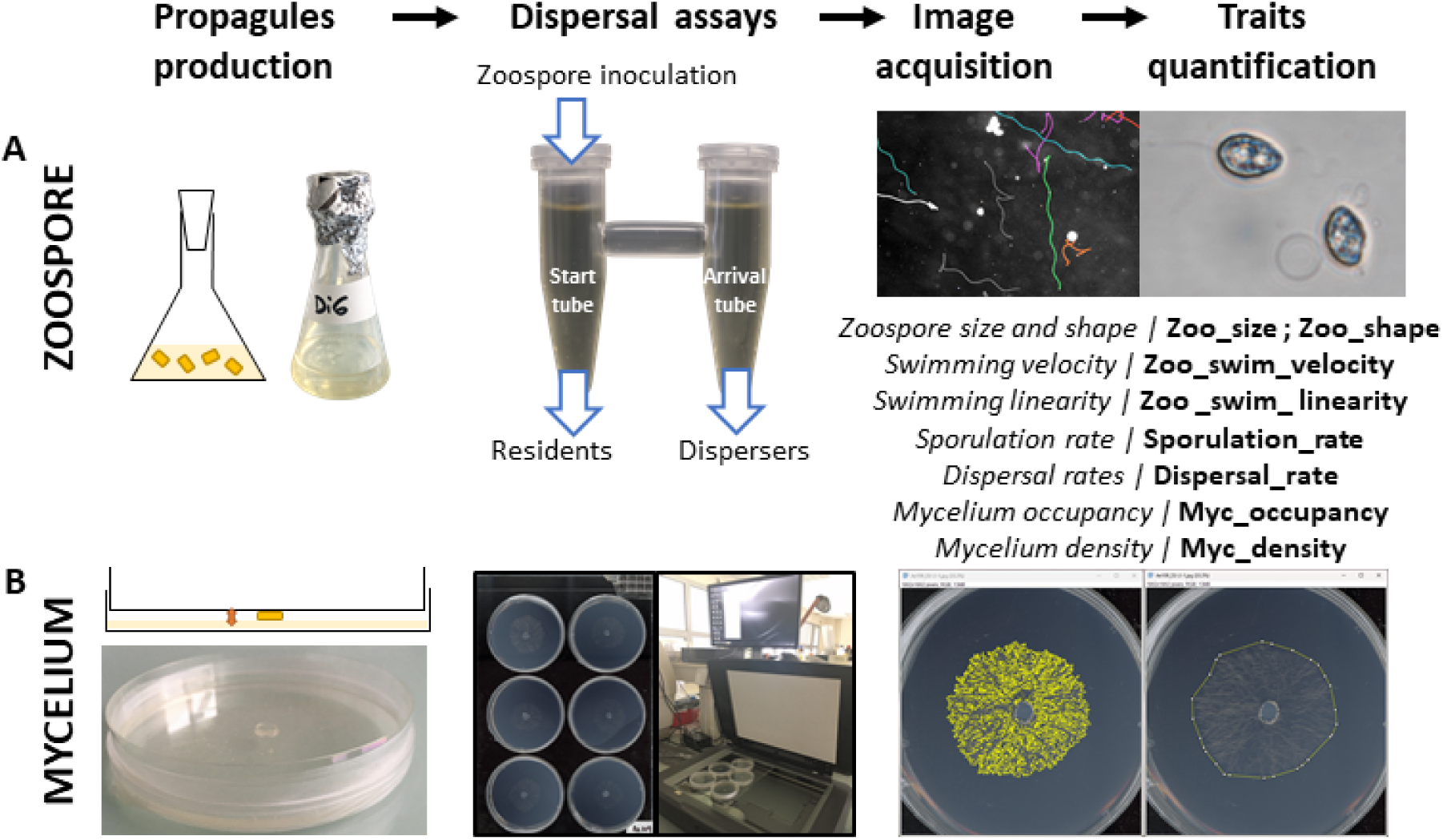
Experimental workflow used to quantify free-living phenotypic traits of *Aphanomyces euteiches*. **(A)** Zoospores produced in liquid culture were introduced into a two-patch dispersal system to quantify zoospore dispersal between connected compartments. Automated image and video analyses were then used to measure zoospore morphological (Zoo_size, Zoo_shape), behavioural (Zoo_swim_velocity, Zoo_swim_linearity), fitness (Sporulation_rate) and dispersal (Dispersal_rate) traits. **(B)** Mycelial colonies grown on CMA medium were automatically imaged and analysed to quantify mycelial development traits, including colony occupancy (Myc_occupancy) and mycelial density (Myc_density). Together, these eight free-living phenotypic traits were used to characterize intraspecific variation and identify the exploration syndrome of *A. euteiches*.

### Whole genome sequencing

#### DNA extraction and long-read sequencing

For each strain, mono-zoosporic isolation was conducted as presented in Camborde *et al*. 2025 with slight modification. Briefly, the number of zoospores was quantified using a Fuchs-Rosenthal slide before dilution in sterile Volvic^®^ water to obtain one zoospore in 10 µL. Around 20 droplets of 10µL were deposited on CMA plates with a distance of 1 to 2 cm and incubated 24h at 22°C in the dark. Single germinated zoospores were isolated and transferred on new CMA plates. Sporulation procedure was again performed for one mono-zoosporic plate per strain, later used for DNA extraction.

High-molecular-weight DNA was obtained from 100 mg of fresh mycelial powder manually ground in mortar and pestle with liquid nitrogen. We extracted DNA with the protocol described in (Mayjonade *et al*. 2016). DNA was quantified using the Qubit dsDNA HS Assay Kit (Life Technologies) and DNA purity was tested on a Nanodrop (Thermo Fisher). Library preparation and sequencing were performed using the SMRTbell prep kit 3.0. Each library was sequenced with the adaptive loading method onto one SMRTcell on the Sequel2 instrument at 100 pM with a 2-h pre-extension and a 30-h movie. Since the strain 201684 did not produce enough biological material to acquire good-quality DNA extraction, we used our previous sequencing results (Kiselev et al. 2022). Due to insufficient genome coverage, Plm1 and EN1-3 strains were excluded from the genomic analysis.

Basecalling was performed using dorado (v1.0.2, (c) 2025 Oxford Nanopore Technologies PLC.) in super-accuracy (sup) duplex mode with the models dna_r10.4.1_e8.2_400bps_sup@v5.2.0 and dna_r10.4.1_e8.2_5khz_stereo@v1.4. Both simplex and duplex reads (corresponding to tags dx:i:0 and dx:i:1 in the BAM file) were assembled using Hifiasm (v0.25.0-r726) (Cheng *et al*. 2024) with the following parameters: --ont --primary --ctg-n 20 -l 0 --n-hap 1. The resulting contigs were filtered to retain only those with a minimum read depth of 15x (based on the rd:i tag in the p_ctg.gfa file). To standardize sequence identifiers across assemblies, the LS1 assembly—which displayed the lowest contig count and the highest contig N50 (4.6 Mb)—was selected as a reference to scaffold the more fragmented assemblies. Synteny identification was carried out using minimap2 (v2.30-r1287) (Li 2021) with the parameter -x asm5, and reference-guided scaffolding was performed using ALLMAPS (v1.5.6) (Tang *et al*. 2015). Mitochondrial genomes were assembled separately using Oatk (v1.0) with the dikarya_mito.fam profile from OatkDB (v20230921) (Zhou *et al*. 2025). Raw sequence reads and assembled genomes are deposited in the NCBI under the BioProject ID PRJNA1522613 and in our in-house data-base, available at: https://lipm-browsers.toulouse.inrae.fr/pub/aphanomics.

#### Whole genome analysis

From the obtained 22 assemblies, whole-genome similarity was estimated using Average Nucleotide Identity (ANI) calculated with FastANI (v1.34) (Jain *et al*. 2018). Genome assemblies in FASTA format were compared in pairwise analyses using default parameters. To assess to genomic relatedness of our sample with five other species of the *Aphanomyces* genus (*A. astaci, A. frigidofilus, A. stellatus, A. laevis, A. cladogamus*), we reconstructed a distance-tree. ANI values were extracted from the tab-separated output generated by FastANI. Clustering with ANI values was visualised using a heatmap. Then, an Analysis of Molecular Variance (AMOVA) was conducted in order to assess how genetic variation is structured among European and North American continents using the *amova* function of the R package pegas (Paradis *et al*. 2025).

#### Infection assays

As the pathotype classification reported in the literature was absent for three strains, harboured intermediate pathotypes for two strains (IN, see above), and was obtained from different experiments conducted at different times (Sup Tab 1), we decided to (re)assess pathogenicity using two complementary methods: *in vivo* and *in vitro* infection assays (Fig S1). Beyond providing indexes of pathogenicity measured for all strains at the same time, our assays have the advantage to quantitatively estimate infectivity.

#### In vivo test

*In vivo* assays were conducted by adapting the method of (Wicker & Rouxel 2001), *i.e.*, the historical method leading to pathotype classification. This laboratory method has the advantage to establish infectivity using several pea lines grown in pots so as to mimic natural conditions as closely as possible. Traditionally, a set of six selected pea lines (*Pisum sativum*), showing similar resistance rankings from fields and controlled conditions assays, is used to assess the variability in virulence of *A. euteiches* (Wicker *et al*. 2003). This set comprises two susceptible lines (Baccara, Capella) and four partially resistant lines showing distinct partial resistance patterns (MN313, 90-2131, 552 and PI180693) (Wicker *et al*. 2003).

Here, a subset of four pea lines that reveal the pathogenicity differences within our collection was used. Firstly, the Baccara line, susceptible to both pathotypes I and III, was used as a control. Then, we used PI180693 and 90-2131 lines, which are resistant to pathotype I but susceptible to pathotype III. Finally, the MN313 line was used as it is resistant to pathotype III but susceptible to pathotype I. Peas were evaluated in controlled condition into a climatic chamber (25°C 14h light / 23°C 10h dark). Five plants were grown in a pot filled with vermiculite substrate. Pots were watered with tap water. Experiment design consisted of four blocks of five plants; for each pea genotype and *A. euteiches* strain, infectivity was measured by calculating the mean disease severity score across the 20 plants evaluated in the four blocks and expressed as a Root Rot Index (RRI). Inoculation consisted in applying an inoculum suspension of 1,000 zoospores per plant on the seven days-old peas (Fig S1A). Seven days post-inoculation, symptoms were visually analysed on roots by extracting plants from the pots. The disease severity score was estimated on each plant, according to the following scale of five scores: 0 = no symptoms; 1 = traces of discoloration on the roots (<25%); 2 = discoloration of 25 to 50% of the roots; 3 = discoloration of 50 to 75% of the roots; 4 = discoloration of >75% of the roots; 5 = plant dead (Fig S1A). An average infection score was calculated for each *A. euteiches* strain by averaging RRI observed within each four blocks across pea genotypes.

#### In vitro test

One limitation of the *in vivo* test is that it does not allow to real-time monitor the root symptoms following infection. Further, a fine-scale inspection of the pathogen impact on each part of the root system development may be difficult to obtain. To overcome these limitations, we adapted an *in vitro* assay developed on the model plant *Medicago truncatula* (Badreddine *et al*. 2008), allowing to observe the root and symptom development in presence of the pathogen.

Four pea lines (*Pisum sativum*) were used: PI180693, 90-2131 and Baccara as in the *in vivo* test, and Caméor (susceptible to both pathotypes I and III) instead of MN313 because we did not have enough material to run the tests on this plant genotype. Pea seeds were surface sterilised with 2.4% active chlorine bleach and were then germinated on petri-dishes containing gelose in the dark at 22°C for one week. Then, four germinated seedlings, each representing a replicate, of the four pea lines were transferred into large petri-dishes (500cm², 245x245x25mm, NUNC) containing M medium (Bécard & Fortin 1988). After 24h, infection was proceeded for each *A. euteiches* strain of the collection on the four seedlings of each pea genotype. Only 22 strains out of 25 produced enough zoospores to proceed to the infection (MF1, NC1 and 22-2 were removed from analysis because of too low sporulation rates). Infection consisted in a 10µL droplet containing 1,000 zoospores deposited 1cm above the root tips. We followed root development into a climatic chamber (22°C 16h light / 20°C 8h dark) for three weeks, but to assess infectivity in the same conditions as the *in vivo* test, we only used data seven days post-inoculation by scanning (Epson) the petri-dishes (Fig S1B). Aerial and root symptoms were analysed visually for each “pea genotype × *A. euteiches* strain” combination. We established a symptom score ranging from 1 to 5 with 1 corresponding to plant without or with light symptoms and 5 to dead plant (Fig S1B). An average infection score was calculated for each *A. euteiches* strain by averaging the symptoms observed across replicates and pea genotypes.

Finally, we used the R function *cor* to test for the significance of correlations (Spearman) between both *in vivo* and *in vitro* infectivity measures to verify that these two methods are consistent.

### Free-living phenotypic traits

Among the 25 strains, MF1 an NC1 (Sup Tab 1), had too low sporulation rates to allow reliable phenotyping and were thus excluded from this part.

### Mycelial traits

Small mycelium plugs (Ø 0.5 cm), cut from the periphery of a seven-day old mycelium produced from a mother culture, were used as inoculum source. In order to limit the aerial mycelial growth and take images in two dimensions, the plugs were positioned on the centre of the petri-dish lid and sealed with the bottom half, so that the two surfaces were nearly in contact, thereby limiting the maximum growth height to ∼0.5 cm (Fig1B). Mycelium were then grown at 22°C in dark in two independent blocks (from two different mother plates) containing each six technical replicates per strain. To follow the mycelial growth, images were taken two times a day (morning and evening) during four days with a flatbed scanner (Epson) at 600 dpi resolution. Images were processed using the ImageJ/Fiji software (Schindelin *et al*. 2012; Schneider *et al*. 2012), which enabled the detection of mycelial hyphae. We then created a polygon around the growth zone by linking the extremities of hyphae in each direction. At each time point, we calculated the polygon perimeter (mm) (Fig1B). As a few scans were unavailable due to technical failure (337 over the 2216 points), these data were processed by linearly interpolating missing values over time for each strain, technical replicate, and independent block using the *na.approx* function of the R package zoo (Zeileis & Grothendieck 2005). A global mycelial development index was then quantified as the trapezoidal integration of the area under the growth curve (*AUC*), obtained by plotting perimeter over time for each technical replicate within blocks. This metric hereafter called *mycelial occupancy* was used as a proxy of global occupancy through time. To obtain a proxy of fine-scale occupancy, we also calculated the *mycelial density* within the polygon at two days (evening measure) of development.

### Zoospore traits

#### Fitness, morphology and behaviour

To obtain a fitness proxy, we measured the sporulation rate of each strain. A subsample of zoospore suspension was quickly vortex to provoke encystment and the number of zoospores per mL was counted under a light microscope using a Fuchs-Rosenthal slide. Sporulation rate was measured on a total of 85 zoospore suspensions (2–5 technical replicates suspensions per strain).

To quantify zoospore morphology and movement-related behaviour, 10 µL of zoospore suspension were placed into chambers of counting slides (Kima precision cell). Videos of 15s were taken under dark-field macroscopy (Axiozoom V16, Zeiss), and analysed with the R video tracking package BEMOVI (Pennekamp *et al*. 2019). Zoospore traits were analysed from 258 videos (6–12 videos per strain from a minimal set of five distinct biological culture). As this package had never been used on *A. euteiches* zoospores before this study, we performed a validation phase by comparing a visual quantification of living and encysted cells under a light microscope with the number of cells detected by Bemovi on 308 zoospore samples comprising all *A. euteiches* strains. We found a high positive and significant correlation (rho = 0.92, p-value < 2.2e-16, Fig S2) validating the methodology.

Four traits were measured with BEMOVI: two morphological traits as *zoospore size* (cell surface area) and *zoospore shape* (ratio of cell major/minor axes, the lower the value, the more rounded the cell is) and two movement traits as *zoospore swimming velocity* (the total distance travelled by the cells divided by the duration of the trajectory) and *zoospore swimming linearity* (ratio between the net Euclidian distance travelled and the total distance travelled, where 1 means a straight line) (Fig1A).

#### Dispersal rate

Dispersal, the movement potentially leading to gene flow (Ronce 2007), is a key eco-evolutionary process influencing the spatial distribution of genotypes whatever the ecosystem, and thus regulates (and is regulated by) species interactions (Bestion *et al*. 2024). To evaluate the dispersal ability of our *A. euteiches* strains, we used two-patch systems, which are commonly used spatially-explicit aquatic microcosms (Altermatt *et al*. 2015; Cayuela *et al*. 2022; Fronhofer *et al*. 2018; Zilio *et al*. 2023). These systems consisted here in two 1.5 mL microtubes connected by a corridor (2.5cm-long silicon pipe with a 4mm internal diameter) filled with sterilized Volvic® water (Fig1A). First, the corridor was closed by a clamp and 300 µL of zoospore suspension was pipetted into one of the two tubes (the ‘start’ tube). After 30min of acclimation, the corridor was opened and zoospores could choose to disperse towards the empty tube (‘arrival’ tube) or stay in the start tube for 2h in the dark at 22°C. Then, the corridor was closed again and 300 µL were pipetted independently in the middle of both start and arrival tubes after three gentle up and down pipetting steps. From these 300µL, 10µL were used to quantify the number of zoospores using BEMOVI (see above). The *dispersal rate* (proxy of dispersal ability) was calculated as the number of dispersing zoospore (found in the arrival tube) divided per the total number of zoospores (zoospores found in the start and arrival tubes). Dispersal rate was analysed from 258 videos (6–12 videos per strain from a minimal set of five distinct biological culture).

### Statistical analyses

All statistics were performed using R studio software version 4.6.0 (R Core Team 2026).

#### Intra- and inter-strains variability

We wanted to assess the variation in free-living phenotypic traits within and between the 23 *A. euteiches* strains, using mycelial traits (Myc_density ; Myc_occupancy), zoospore morphological traits (Zoo_size ; Zoo_shape), zoospore behavioural traits (Zoo_swim_velocity ; Zoo_swim_linearity), two fitness-related trait (Sporulation_rate ; Dispersal_rate). Firstly, we computed basic statistical parameters such as mean, median, standard deviation and variance at both overall collection and strain levels. Then, the contributions of the strain effect to the total variability of mean free-living phenotypic traits were quantified using repeatability estimation based on LMMs fitted by restricted maximum likelihood (REML). Analyses were performed using the *rptGaussian* function of the rptR package (Stoffel *et al*. 2017), assuming a Gaussian data distribution. Repeatability estimates were derived with 1,000 bootstrap iterations to assess uncertainty. Finally, to determine which and how many strains differed from the reference strain RB84 for each phenotypic trait, we fitted LMMs using REML (via the *lmer* function of the lme4 package, Bates *et al*. 2015), with strain treated as a fixed effect and the replicate as a random effect.

#### Existence of trait syndrome in A. *euteiches* and hierarchical clustering of the collection

To test for the existence of trait syndrome, we first calculated spearman correlations between all free-living phenotypic trait pairs (R function *cor*). Then, in order to assess potential links between the way infectivity is measured and phenotypic variables, we calculated spearman correlations between these phenotypic traits and infectivity *in vivo* and *in vitro* measures. Secondly, as some traits were correlated, a Principal Component Analysis (PCA) was used to obtain axes of phenotypic variation structuring the species diversity using the *prcomp* function of the stats package. For each free-living phenotypic trait and strain, we calculated a standardized estimation of the deviation of a given strain phenotype to the mean overall strains using a Z-score calculated as follows: Z-score = (X – X̄) / σ where X is a trait mean for a strain, X□ is the trait mean overall strains and σ is the trait standard deviation overall strains. These Z-scored were then plotted as a heatmap (package pheatmap, function *pheatmap*) (Kolde 2025). A hierarchical clustering was done in order to group strains based on Euclidian distances with the ward.D2 method. The robustness of this hierarchical clustering was evaluated with bootstrap including 1,000 iterations. Approximately Unbiased (AU) *p*-values and Bootstrap Probability (BP) values were measured for each cluster node using the *pvclust* function of the pvclust package (Ryota *et al*. 2019). Finally, these clusters were visualized on the PCA to confront them to the main detected axes of phenotypic variation. Phenotypic trait profiles specificity according to each cluster was tested with a Kruskal-Wallis test.

#### Linking phenotypic variability to the genomic diversity, geographic origin and pathogenicity of strains

We wanted to assess if there was a sub-structure in the described phenotypic diversity beyond a general phenotypic syndrome according to three scales: strains’ genomic diversity (North American *vs.* European strains clusters, see ANI results), strains’ geographic origin (Denmark, France, Finland, Sweden, USA) and strains’ pathogenicity (as traditional pathotype or with indexes measured with *in vivo* and *in vitro* tests). We first estimated the repeatability of trait values according to these three scales with the rpt package as presented before (see Intra- and inter-strain variability). Then, to determine if groups of strains based on these three scales situate in similar spaces of the described phenotypic syndromes, we used a Permutational Multivariate Analysis of Variance (PERMANOVA) to test for significant differences in the centroids of PCA coordinates from the first four dimensions among strains groups (R package vegan, function *adonis2* with Euclidian distance and 9999 permutations) (Oksanen 2025). Finally, Kruskal–Wallis tests were used to compare levels of strains’ pathogenicity (pathotype*, in vivo* and *in vitro* infectivity tests) among the phenotypic clusters.

## RESULTS

### Genetic structure in *A. euteiches*

Whole-genome assemblies of 22 strains of our *A. euteiches* collection were obtained to study their genetic diversity and structure.

An average of 5.3 Gb (range: 3.1–9.7 Gb) of sequencing data was generated per sample, corresponding to a sequencing depth between 44× and 139× (mean: 76×). Read N50 values ranged from 14.4 to 26.4 kb, which enabled the generation of highly contiguous assemblies with at most 5 gaps. The resulting assemblies showed scaffold N50 values between 4.0 and 4.6 Mb and total genome sizes ranging from 68.4 to 70.4 Mb (Sup Tab 2).

The distance-tree including five other *Aphanomyces* species showed a marked genetic distance between our strains and these species, and revealed two main genetic groups (Fig S3A). Pairwise ANI exceeded 98% across all strains indicating limited genetic diversity at the collection level. Nevertheless, ANI analyses confirmed the existence of two genetically-distinct groups (yellow and purple colours in Fig 2). This was validated by the AMOVA results as the continent (Europe *vs.* North America) explained 99.7% of the total genetic variance with very low within-group variation (Φ = 0.997, p < 0.001). This low within-continent variation is particularly obvious within the Burgundy (France) sub-cluster (mean ANI = 99.998% ± 0.026%, Fig S3B). Notably, this sub-cluster comprises strains with the intermediate (IN) pathotype, distinguishing it from other European isolates (Fig 2 & S3). Altogether, these analyses revealed a geographically structured strain collection despite an overall low level of genomic diversity.

**Figure 2.**
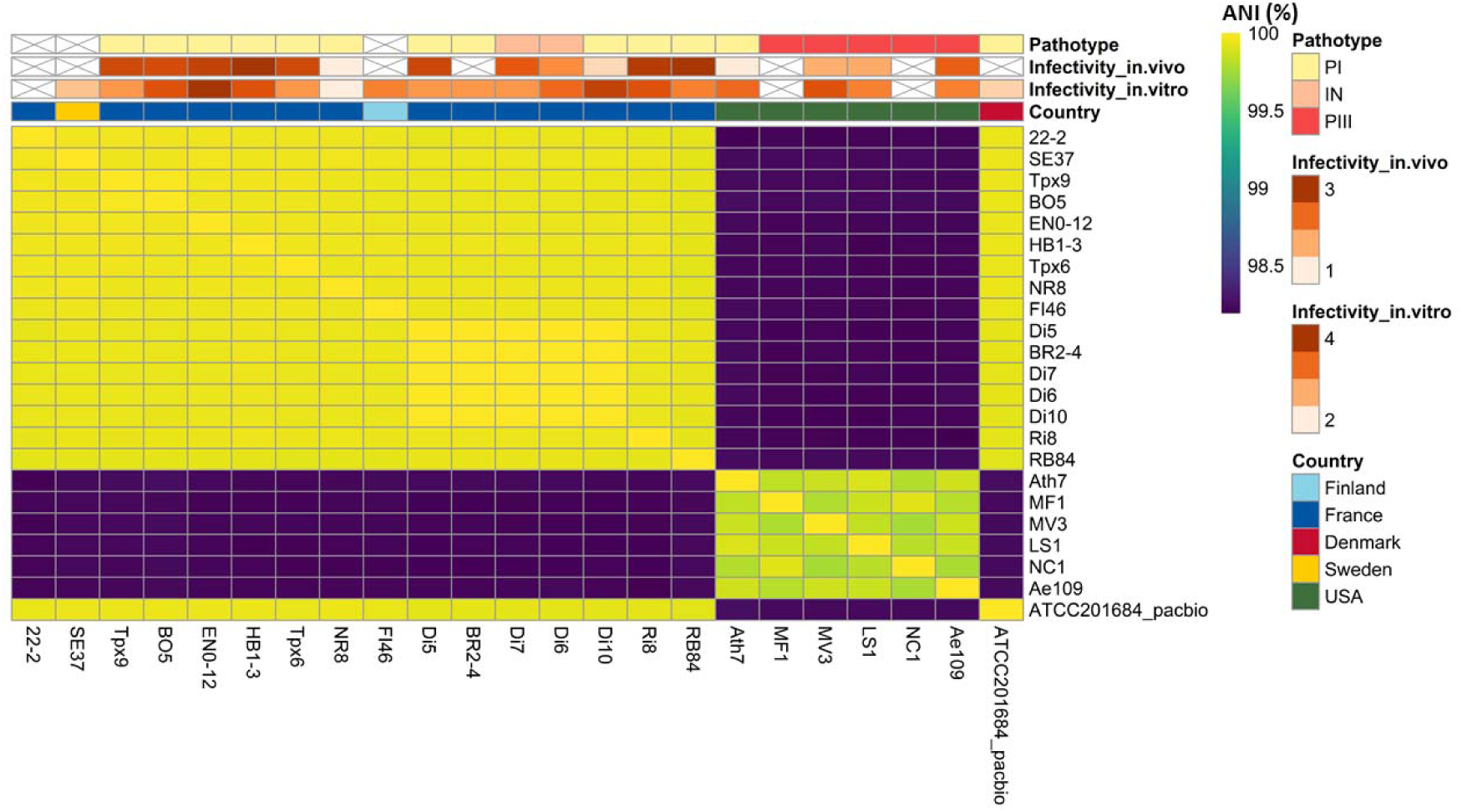
Whole-genome Average Nucleotide Identity (ANI) of the Aphanomyces euteiches strain collection. Pairwise ANI values were calculated among the 23 sequenced strains and visualized as a heatmap, with colours ranging from dark purple (lower ANI) to bright yellow (higher ANI). The heatmap reveals two major genetic groups corresponding to European and North American strains, with high genomic similarity within European strains and greater genetic diversity among North American strains. Annotation bars indicate pathotype, infectivity measured *in vitro* and *in vivo*, and country of origin. Cross indicated absence of data.

### Trait variation in *A. euteiches*

We phenotyped our *A. euteiches* collection for eight free-living phenotypic traits. From this phenotyping of mycelium and zoospores, we analysed variation both at the collection level (Sup Tab 3) and at the strain level (Sup Tab 4, Sup Tab 5, Fig S4), notably through repeatability estimates (Tab 1).

**Table 1.** Repeatability estimates of the eight free-living phenotypic traits according to strain, geographic origin, genetic group, pathotype, and infectivity measured using the *in vitro* and *in vivo* assays. Repeatability (R) was estimated using Gaussian linear mixed models fitted by restricted maximum likelihood (REML) with the rptGaussian function of the *rptR* package. Standard errors (SE), 95% confidence intervals (95% CI) and associated *P*-values are reported for each estimate and written in bold when significant.

| Trait | Strain |  |  |  | Geography origin (country of isolation) |  |  |  | Genetic group (Europe vs USA) |  |  |  |
| --- | --- | --- | --- | --- | --- | --- | --- | --- | --- | --- | --- | --- |
|  | R | SE | CI | P | R | SE | CI | P | R | SE | CI | P |
| *Myc_occupancy* | 0.69<br>6 | 0.10<br>4 | [0.437,<br>0.828] | **6.76e-29** | 0 | 0.02<br>4 | [0, 0.089] | 1 | 0 | 0.02<br>5 | [0, 0.085] | 1 |
| *Myc_density* | 0.66<br>5 | 0.07<br>8 | [0.478,<br>0.783] | **1.4e-37** | 0.55<br>8 | 0.18<br>6 | [0.099,<br>0.795] | **2.79e-18** | 0.42<br>8 | 0.24<br>9 | [0, 0.796] | **3.34e-12** |
| *Zoo_size* | 0.21<br>1 | 0.06<br>7 | [0.081,<br>0.336] | **6.69e-08** | 0.18<br>9 | 0.11<br>8 | [0, 0.42] | **1.81e-09** | 0.32<br>9 | 0.22<br>1 | [0, 0.717] | **6.68e-09** |
| *Zoo_shape* | 0.02<br>1 | 0.02<br>7 | [0, 0.093] | 0.353 | 0 | 0.02<br>2 | [0, 0.075] | 1 | 0 | 0.01<br>1 | [0, 0.041] | 1 |
| *Zoo_swim_velocity* | 0.38<br>2 | 0.08<br>7 | [0.201,<br>0.534] | **1.67e-16** | 0.16<br>8 | 0.11<br>4 | [0, 0.412] | **0.00172** | 0.05<br>9 | 0.07<br>5 | [0, 0.254] | 0.0949 |
| *Zoo_swim_linearity* | 0.50<br>9 | 0.08<br>4 | [0.323,<br>0.642] | **3.68e-27** | 0.26<br>4 | 0.14<br>5 | [0, 0.538] | **0.00408** | 0.06<br>9 | 0.08 | [0, 0.283] | 0.0596 |
| *Sporulation_rate* | 0.63<br>7 | 0.09<br>8 | [0.407,<br>0.788] | **2.4e-10** | 0.33<br>6 | 0.20<br>1 | [0, 0.681] | **0.000545** | 0.33<br>3 | 0.23<br>5 | [0, 0.736] | **0.00202** |
| *Dispersal_rate* | 0.01 | 0.02<br>7 | [0, 0.091] | 0.479 | 0.13<br>6 | 0.14<br>7 | [0, 0.523] | 0.0669 | 0.13<br>6 | 0.13<br>4 | [0, 0.464] | 0.0669 |

| Trait | Pathotype |  |  |  | Infectivity in vitro |  |  |  | Infectivity in vivo |  |  |  |
| --- | --- | --- | --- | --- | --- | --- | --- | --- | --- | --- | --- | --- |
|  | R | SE | CI | P | R | SE | CI | P | R | SE | CI | P |
| *Myc_occupancy* | 0.02 | 0.03<br>7 | [0, 0.126] | 0.375 | 0.80<br>2 | 0.09<br>4 | [0.552, 0.9] | **7.75e-50** | 0.64<br>3 | 0.09 | [0.41,<br>0.769] | **9.98e-37** |
| *Myc_density* | 0.09<br>6 | 0.10<br>2 | [0, 0.35] | **0.00926** | 0.48<br>1 | 0.14<br>2 | [0.156,<br>0.681] | **9.62e-12** | 0.52<br>4 | 0.10<br>4 | [0.269,<br>0.675] | **2.01e-18** |
| *Zoo_size* | 0.28<br>7 | 0.17 | [0, 0.589] | **5.24e-06** | 0.10<br>5 | 0.06<br>7 | [0, 0.256] | **0.000119** | 0.25<br>8 | 0.08<br>3 | [0.091,<br>0.411] | **1.55e-08** |
| *Zoo_shape* | 0 | 0.01<br>7 | [0, 0.064] | 0.5 | 0.00<br>6 | 0.01<br>7 | [0, 0.055] | 0.415 | 0.04<br>8 | 0.03<br>9 | [0, 0.136] | 0.11 |
| *Zoo_swim_velocity* | 0.01<br>3 | 0.02<br>8 | [0, 0.099] | 1 | 0.11<br>3 | 0.06<br>8 | [0, 0.275] | **0.00221** | 0.29<br>2 | 0.08<br>7 | [0.12,<br>0.464] | **2.24e-10** |
| *Zoo_swim_linearity* | 0.2 | 0.14<br>6 | [0, 0.512] | **0.00512** | 0.22<br>2 | 0.1 | [0.045,<br>0.423] | **5.35e-06** | 0.50<br>9 | 0.09<br>6 | [0.274,<br>0.666] | **1.11e-23** |
| *Sporulation_rate* | 0.64<br>5 | 0.25<br>6 | [0, 0.881] | **8.25e-06** | 0.25<br>2 | 0.12<br>8 | [0, 0.48] | **0.00252** | 0.62<br>7 | 0.11<br>3 | [0.367,<br>0.786] | **5.21e-09** |
| *Dispersal_rate* | 0.08<br>1 | 0.08 | [0, 0.288] | **0.0277** | 0.02<br>6 | 0.03<br>7 | [0, 0.133] | 0.226 | 0 | 0.01<br>8 | [0, 0.06] | 1 |

### Mycelial growth strategies

*Myc_density* showed high variability (CV = 0.456) while *Myc_occupancy* was less variable (CV = 0.124, Sup Tab 3), yet repeatability estimates were high and comparable for the two traits (*Myc_occupancy* R= 0.696, p-value = 6.76e^-29^ and *Myc_density* R=0.665, p-value = 1.4e^-37^, Tab 1). This indicates a strong effect of strain identity on the expression of mycelial traits, with a greater heterogeneity between strains in local than in global mycelial occupancy. Inspecting these strain-specific patterns revealed the existence of contrasting mycelial growth phenotypes within the collection. For instance, strains 201684 and 22-2 showed low biomass accumulation both globally and locally (*Myc_occupancy:* 259 ± 49 and 285 ± 22, respectively, and *Myc_density:* 186 ± 68 and 269 ± 56, respectively). In contrast, strain EN0-12, BO5, Ae109, and EN1-3 showed dense and extensive mycelial development (*Myc_occupancy* >460, *Myc_density* >600) (Sup Tab 4A and Fig S4).

Overall, mycelial traits exhibited substantial strain-dependent variation, supporting the existence of contrasting mycelial growth phenotypes within the collection.

### Zoospore morphology and behaviour

Zoospore behavioural traits showed higher variability than zoospore morphological traits (CV = 0.184 and CV = 0.202 respectively for *Zoo_swim_velocity* and *Zoo_swim_linearity* compared to CV = 0.154 and CV = 0.104, respectively for *Zoo_size* and *Zoo_shape*, Sup Tab 3), suggesting stronger constraints on morphology than on movement. Accordingly, repeatability estimates were higher for behavioural traits (*Zoo_swim_linearity* R=0.509, p-value = 3.68e^-27^ and *Zoo_swim_velocity* R=0.382, p-value = 1.67e^-16^) than for morphological traits (*Zoo_size* R=0.211, p-value = 6.69e^-8^, Tab 1). Among our strains, Ae109, Plm1, Ath7 or LS1 produced larger zoospores (>350) with a rounder shape (<1.33), while others, such as Tpx6, Di 5, Di7 or RB84 produced smaller zoospores (<296) with a more elongated shape (>1.35) (Sup Tab 5A; Fig S4). Regarding movement, some strains had faster-moving zoospores with straighter trajectories such as for Di7, 201684, EN1-3 and EN0-12 strains (Sup Tab 5A; Fig S4).

Therefore, like for mycelial traits, zoospore morphological and behavioural traits exhibited strain-dependent variation, but behavioural traits were more variable and strain-specific than morphological traits.

### Sporulation and dispersal rates

*Sporulation_rate* was the most variable trait across strains at the collection scale (CV = 0.85; from 1,250 to 817,500 zoospores per mL, Sup Tab 3) with very high repeatability estimates (R=0.637, p-value = 2.4e^-10^, Tab 1). The most productive strains generated nearly 22-fold more zoospores than the least sporulating strains (strains Di5–Di7 produced ∼520,000 zoospores, strains Plm1, SE37, and LS1 produced less than 56,000 spores (Sup Tab 5A; Fig S4). By contrast, repeatability was low and non-significant for *Dispersal_rate* (R=0.01, p-value = 0.479, Tab 1), which was generally very low (average = 0.029 ± 0.07, from 0 to 0.5, Tab 2). In other words, dispersal events were infrequent in our two-patch systems (Sup Tab 5A). However, strains LS1, MV3, FI46 and SE37 exhibited higher mean dispersal rate (>0.05) and greater variability (SD = 0.168 and 0.167, for LS1 and SE37 respectively), suggesting occasional dispersal pulses.

**Table 2.** Summary of the phenotypic characteristics of the six clusters identified from the eight free-living phenotypic traits. . For each cluster, the predominant characteristics of mycelial traits, zoospore morphological and behavioural traits, zoospore fitness and dispersal traits are summarized. Robust clusters supported by the hierarchical clustering analysis (AU ≥ 95%) are highlighted in grey. Specific traits according to significative differences between clusters (Kruskal-Wallis tests, Sup Tab 8) are written in bold.

| Cluster ID | Strains (Origin) | Mycelial traits<br>*occupancy + density* | Zoospore morphological traits<br>*size + shape* | Zoospore behavioural traits<br>*velocity + linearity* | Fitness-related trait<br>*sporulation* | Dispersal trait<br>*dispersal* |
| --- | --- | --- | --- | --- | --- | --- |
| 1 | BR2-4 (FR) | Moderately sparse mycelial network | Large, elongated zoospores | **Slowest swimming velocity** with tortuous trajectories | Low sporulation rate | Low dispersal rate |
| 2 | MV3, LS1, Plm1, Ath7, Ae109 (USA) \| SE37 (SE) \| HB1-3, NR8 (FR) | Extensive mycelial network | **Larger and rounder zoospores** | Slow swimming with moderately tortuous trajectories | Low sporulation rate | Contrasting dispersal rates |
| 3 | 201684 (DE) \| 22-2, Tpx6 (FR) | **Sparse mycelial network** | Small, elongated zoospores | Contrasting swimming behaviours | Contrasting sporulation rates | Low dispersal rate |
| 4 | FI46 (FI) \| Tpx9, RB84 (FR) | Moderately sparse but extensive mycelial network | Small, elongated zoospores | Predominantly tortuous swimming | Moderately high sporulation rate | Contrasting dispersal rates |
| 5 | Di5, Di6, Di7, Di10 (FR) | Moderately dense and extensive mycelial network | Small and elongated zoospore | Fast swimming with **highly linear trajectories** | High sporulation rate | Low to moderate dispersal rate |
| 6 | EN1-3, Ri8, BO5, EN0-12 (FR) | Dense and **extensive mycelial network** | Small, elongated zoospores | Fast swimming with linear trajectories | Intermediate sporulation rate | Low to moderate dispersal rate |

Overall, our fitness proxy was strongly structured at the strain level, whereas dispersal remained highly variable within strains and showed little differentiation among strains.

### Comparison with the reference RB84 strain

For almost all traits, the majority of strains had phenotypic values significantly different from the reference strain RB84 (Sup Tab 6). It was only for *Zoo_shape* (at the exception of three strains) and *Dispersal_rate* that RB84 had similar values than the other strains.

### ‘Exploration’ syndrome in *A. euteiches* and strains clustering

We wanted to know if the measured intraspecific variation was globally organised into a phenotypic syndrome underlying the exploration capacity of the species. We first examined correlations between traits at the collection scale (Fig S5) and then studied the syndrome with a PCA (Fig 3). To explore whether trait variation reflects broader ecological or adaptive patterns, we performed hierarchical clustering of strains based on the Z-score of the eight free-living phenotypic traits (Fig 4) and we compare traits between the obtained clusters with Kruskal-Wallis tests to highlight cluster phenotypic specificities.

**Figure 3.**
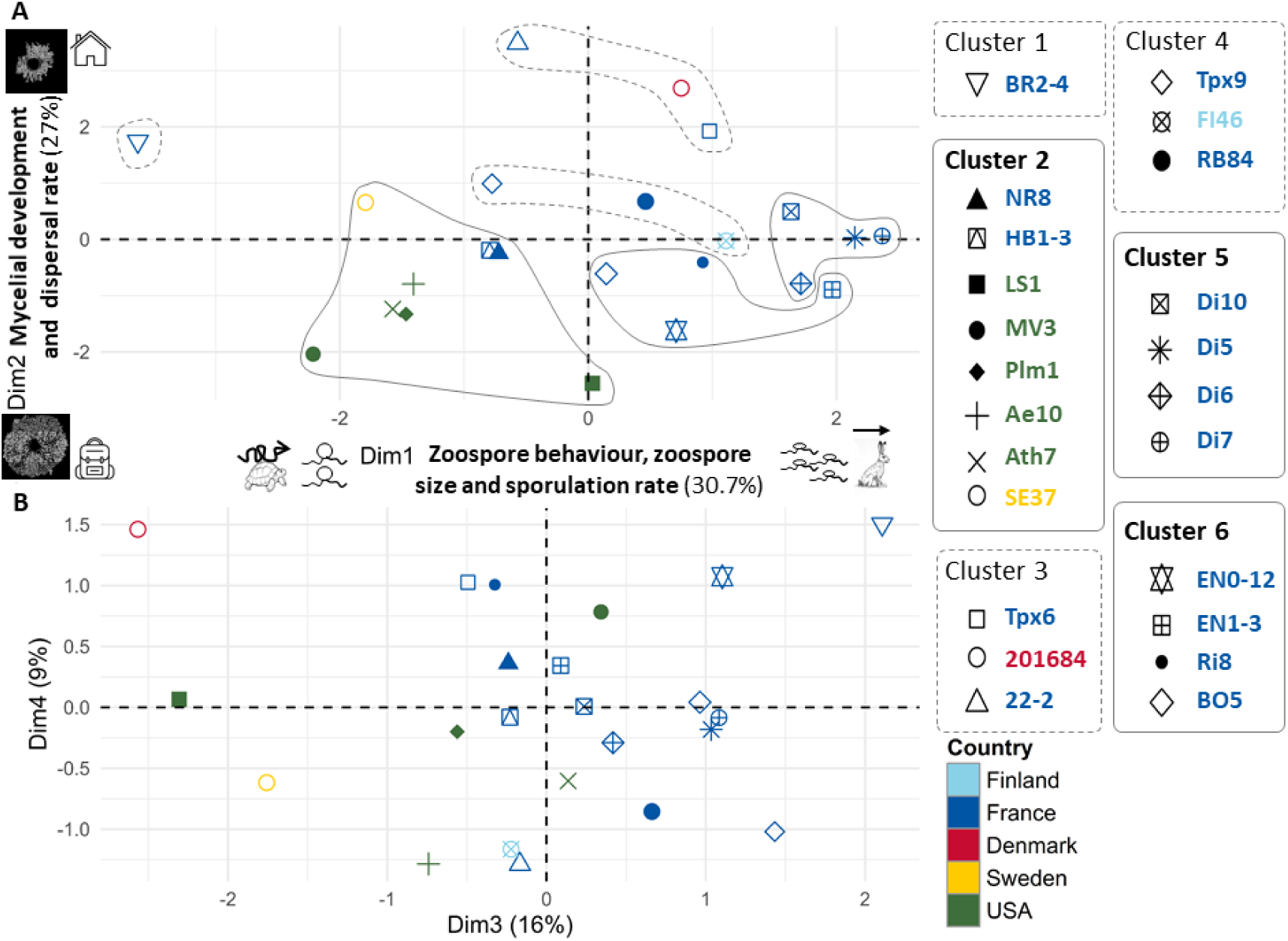
Principal component analysis of free-living phenotypic traits in the *A. euteiches* collection. PCA was performed using the mean values of the eight phenotypic traits measured for the 23 strains. The percentage of variance explained by each dimension is indicated on the corresponding axes. Symbols represent individual strains and colours indicate the country of origin. **(A)** PCA of Dimensions 1 and 2. Dimension 1 is mainly associated with zoospore behaviour, zoospore size and sporulation rate, whereas Dimension 2 is associated with mycelial development (occupancy and density) and zoospore dispersal. Icons illustrate the ecological interpretation of the two dimensions and reveal an ‘exploration’ syndrome in *A. euteiches*. On the first dimension: high production of small and elongated zoospore having a fast and linear swimming behavior (rabit and zoospore symbols) *vs.* low production of big and round zoospore having a slow and tortuous swimming behavior (turtle and zoospore symbols). On the second dimension: high exploration through dense and extensive mycelium and dispersive zoospore (extensive mycelial and backpack symbols) *vs.* low exploration with sparse and localized mycelium and non-dispersive zoospore (small mycelial and house symbols). The six clusters are represented in bold and grey lines (robustly supported) and in dashed lines (not supported). **(B)** PCA of Dimensions 3 and 4.

**Figure 4.**
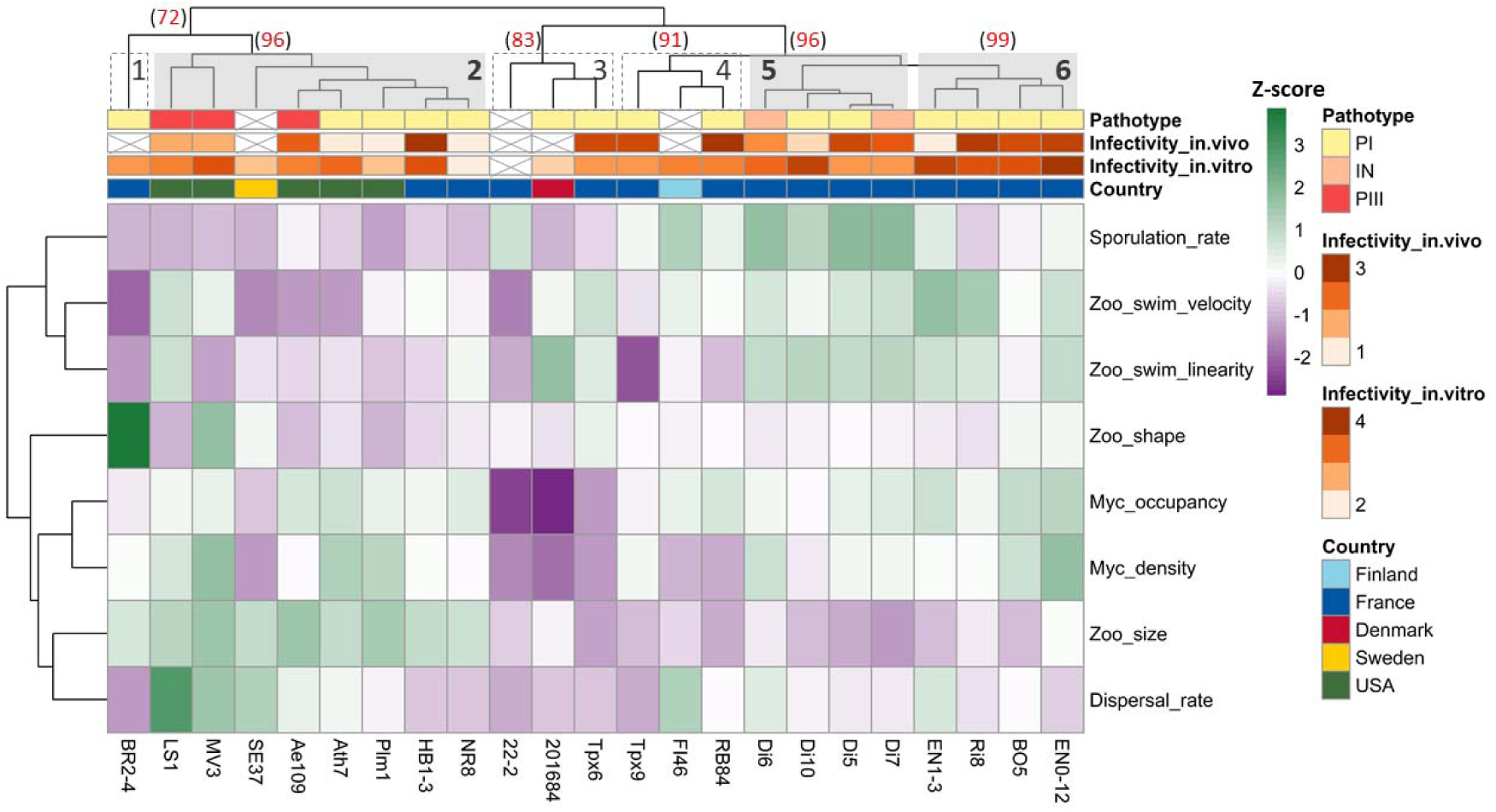
Hierarchical clustering of *Aphanomyces* euteiches strains based on free-living phenotypic traits. Heatmap showing Z-score-normalized values of the eight phenotypic traits measured across the strain collection. Hierarchical clustering was performed using Euclidean distance and Ward D2 linkage. Annotation bars indicate pathotype, infectivity measured *in vitro* and *in vivo*, and country of origin. Cross indicated absence of data. Clusters supported by Approximately Unbiased (AU added to the dendrogram in red) values ≥95% are indicated in bold by solid grey boxes, whereas less supported clusters (AU <95%) are delimited by dashed boxes.

### Correlation between pairs of traits

No significant pairwise correlation was found between mycelial and zoospore traits (see nonetheless the marginal correlation between *Myc_density* and *Zoo_size*), while significant correlations were observed between mycelial traits and between zoospore traits respectively (Fig S5). *Myc_occupancy* and *Myc_density* were positively correlated (r= 0.55, p=7.87e^-5^), indicating that strains with high radial mycelial expansion developed dense networks (Fig S5). For the two zoospore morphological traits, no significant correlation was observed. In contrast, *Zoo_swim_linearity* and *Zoo_swim_velocity* were positively correlated (r = 0.66, p= 1.87e^-3^), indicating that faster-moving zoospores had straighter trajectories (Fig S5). However, behavioural traits did not align with morphological traits, at the exception of a marginal correlation between *Zoo_size* and *Zoo_swim_velocity* (Fig S5). *Sporulation_rate* was negatively correlated with *Zoo_size* (r = –0.70, p= 3.64e^-4^), indicating that strains producing smaller zoospores also exhibited higher asexual reproductive output (Fig S5). Noteworthy, *Dispersal_rate* was not significantly correlated to any other trait, but nonetheless marginally related with *Zoo_size*.

Together, these analyses reveal that some free-living phenotypic traits are correlated within, but not strongly between, the mycelial and zoospore stages of *A. euteiches*.

### PCA ordination and clustering

To better explore correlations, we performed a PCA. Its first four dimensions explained 83% of the total variance and thus captured the majority of the covariation pattern among our measured free-living phenotypic traits (Fig 3; Fig S6; Sup Tab 7A). Dimension 1 was associated with zoospore morphological, behavioural and fitness traits while Dimension 2 was associated with mycelial development and zoospore dispersal. Positively associated with Dimension 1, *Zoo_swim_velocity*, *Zoo_swim_linearity* and *Sporulation_rate*, together, contribute for more than 60 % of this axis variance, whereas *Zoo_size* and *Zoo_shape*, negatively associated with it were contributing for around 30 % of its variance (Sup Tab 7B and C). Dimension 2 was negatively associated with the two mycelial traits, contributing together for more than 60 % of this axis variance and *Dispersal_rate* contributing to 18 % (Sup Tab 7B and C). These two first dimension summarise the ‘exploration’ syndrome in *A. euteiches*; zoospore with rapid and linear swimming are also small, numerous and elongated (Dim 1) and mycelium with high development (expansion and density) produce zoospore with higher dispersal rate (Dim 2).

The hierarchical clustering revealed six clusters of one to eight strains. Three (Clusters 2, 5 and 6) were robustly supported (AU p-values > 95%, Fig S7), each associated with distinct trait profiles and geographic origins (Fig 4). Associated key trait profiles are presented in Tab 2 and supported by significant traits differentiation (Sup Tab 8). Cluster 2, which includes all North American strains, was mainly characterised by significantly rounder and larger zoospores (Sup Tab 8) whereas Cluster 5, composed of strains from the Burgundy area of France was characterized by a high production of smaller and more elongated zoospores with fast and highly linear swimming behaviour (Sup Tab 8). Cluster 6, composed exclusively of French strains, showed the highest mycelial development whereas Cluster 3 was characterized by a reduced mycelial development (Sup Tab 8). Clear separations of these four clusters appeared respectively along the PCA Dimension 1 for Cluster 2 and 5 and Dimension 2 for Cluster 6 and 3 (Fig 3).

Altogether, these analyses revealed that variation of free-living phenotypic traits in *A. euteiches* draw distinct strain clusters associated with specific traits profiles alongside with the emergence of an ‘exploration’ syndrome at the species scale.

### Impact of geography, genomic and infectivity on phenotypic variation

We next investigated if geographical origin (five countries of isolation), genetic group (Europe *vs.* North America) or pathogenicity could explain of the observed phenotypic variation. We estimated the repeatability for each trait values at these three scales and used PERMANOVA to quantify their contribution to the described phenotypic clustering.

### Geographic origin & Genetic group

*Myc_density*, *Sporulation_rate* and *Zoo_size* showed high repeatability estimates according to geographical origin (R = 0.558 p-value = 2.79e^-18^, R = 0.336 p-value = 0.000545, R = 0.189 p-value = 1.81e^-09^) and genetic groups (R = 0.428 p-value = 3.34e^-12^, R = 0.333 p-value = 0.00202, R = 0.329 p-value = 6.68e^-09^). In addition, the two zoospore behavioural traits showed significant repeatability across geographic origin, yet less (Tab 1).

PERMANOVA confirmed that both geographical origin and genetic group, significantly influenced the described phenotypic clustering by explaining, respectively, 38% (p-value = 0.0083) and 27% (p-value = 0.0003) of the variance in Dimensions 1 and 2 respectively (Sup Tab 9).

These results show that geography and genomic variability influence most of our phenotypic traits and contribute significantly to the strain clustering we described before.

### Pathogenicity

For pathogenicity we used the general pathotype classification reported in the literature (PI, IN or PIII) and our quantitative estimations based on an *in vivo* infectivity assay, commonly used in literature to assess pathogenicity on pea (Wicker & Rouxel 2001), and an *in vitro* infectivity assay, primarily designed on the model plant *M. truncatula* (Badreddine *et al*. 2008).

No significant correlation was observed between the two infectivity assays measures (Fig S8). We then examined correlations between infectivity metrics and free-living phenotypic traits, only *in vitro* infectivity significantly correlated positively with mycelial traits, but notice that it was also marginally correlated with *Zoo_swim_velocity* (Fig S5). Interestingly, four phenotypic traits (*Myc_density*, *Zoo_size*, *Zoo_swim_linearity* and *Sporulation_rate*) showed a significant effect of infectivity estimations on repeatability regardless of how pathogenicity was evaluated, with the highest repeatabilities generally obtained for *in vivo* assay (Tab 1). In contrast, *Myc_occupancy* and *Zoo_swim_velocity* repeatability were only associated with *in vitro* and *in vivo* infectivity assays and *Dispersal_rate* exclusively with the general pathotype classification.

The PERMANOVAs showed that among the pathogenicity levels, only the general pathotype explained significantly the variance (26%) in PCA Dimensions 1 and 2 (p = 0.0275) (Sup Tab 9). However, a relationship close to significance between the *in vitro* infectivity measures and strain position along Dimensions 1 and 2 also appears (11%, p = 0.0884). These relationships show a link between the pathogenicity of the strains and their measures of traits associated with the exploration syndrome. Finally, no significant differences in all three pathogenicity levels was found between strain clustering (Sup Tab 8A).

Taken together, these analyses indicate that pathogen infectivity contribute to the phenotypic structuring of the strain collection, while they influence different components of the exploration syndrome.

## DISCUSSION

Free-living stages of oomycetes, and more generally of soil-borne pathogens, are often overlooked in the study of plant-pathogen interactions. This study provides evidence for the existence of high intraspecific variation in free-living traits of *A. euteiches*, further structured in an exploration syndrome describing some ecological specialization of this plant pathogen. We also detected phenotypic clusters along the two major axis of the exploration syndrome, whose variance is partially explained by geographic origin, genomic diversity and strain pathogenicity. Interestingly, the *A. euteiches* strains collection revealed that most genetic variation was partitioned between European and North American continents with limited variation within each geographic subgroup, and that it is possible to link infectivity with particular phenotypic values of free-living stages.

Whole-genome sequencing generated assemblies with genome sizes consistent with the reference genome of *A. euteiches* strain ATCC201684 (72 Mb; (Kiselev *et al*. 2022)). Our work also confirmed previous reports of low genetic diversity in *A. euteiches* (Grünwald & Hoheisel 2006; Malvick & Percich 1998; Mieuzet *et al*. 2016; Quillévéré-Hamard *et al*. 2018; Wicker & Rouxel 2001). However, this apparent genomic homogeneity masked a pronounced geographic structure, with near-complete genetic differentiation between European and North American strains. This pattern is consistent with results reported by Le May et al (2018), who identified two different genetic clusters of French and North American isolates geographically separated, based on SRAP (Sequence Related Amplified Polymorphisms) markers. Geographic isolation plays a major role in shaping the genetic structure of the species, whereas pathotype differentiation is more likely explained by localized adaptive changes than by genome-wide divergence. A hypothesis could be that the high genetic homogeneity of European strains, particularly within the Burgundy sub-cluster, may reflect a founder effect following the introduction of *A. euteiches* into Europe from a restricted North American source, however this need further investigation. Subsequent clonal propagation through asexual reproduction, together with long-term survival of oospores in soil and the predominantly homothallic reproductive mode of the pathogen (Facon *et al*. 2006; Grünwald & Hoheisel 2006), may have further reduced genetic diversity. By contrast, the greater genetic divergence observed among North American strains likely reflects a longer evolutionary history and host-associated differentiation driven by agricultural practices or host specialization, particularly between alfalfa (MF1, NC1) and pea isolates (Malvick *et al*. 2009; Moussart *et al*. 2024). Nevertheless, experimental evidence of outcrossing in controlled conditions (Shang *et al*. 2000) and the high heterozygosity reported in some French populations (Kälin *et al*. 2026; Quillévéré-Hamard *et al*. 2018) suggest that sexual recombination may occur in the considered homothallic *A. euteiches*, which could increase genetic diversity. Alternatively, the marked geographic structure observed here, combined with the worldwide distribution and broad host range *(pea, alfalfa, vetch faba bean, bean and lentil)* of *A. euteiches* (Malvick *et al*. 2009; Moussart *et al*. 2008; Tivoli *et al*. 2006) may also be consistent with the existence of two well-separated lineages from two continents, whose taxonomic status might be to clarified.

Trait-based approaches are increasingly being considered to predict ecological strategies in fungi and oomycetes, although they remain relatively uncommon because trait definition and measurement are conceptually and methodologically challenging. In plant pathology, quantitative traits are traditionally assessed during host colonization, based on the assumption that microbial pathogenicity, dispersal, and survival emerge from interactions between the pathogen, the host, and the environment (Delmas *et al*. 2014; Delmotte *et al*. 2014; Lannou 2012). Consequently, traits such as lesion size, sporulation rate, and latent period are generally measured manually, limiting experimental throughput. Most studies on phytopathogenic oomycetes have therefore focused on host-associated traits or on zoospore responses to host-derived cues, particularly chemotaxis towards root exudates (Cohen *et al*. 2025; Kasteel *et al*. 2023, 2024; Lupatelli *et al*. 2023). However, evidence suggests that zoospore motility alone may enhance pathogen fitness by promoting exploration and dispersal through the surrounding environment (Blanco & Judelson 2005; Carlile 1983; Sujkowski *et al*. 2000). Yet, these studies have generally examined free-living traits individually and in a limited number of strains, precluding an integrated view of phenotypic variation at the species level. Here, we combined an image- and video-based phenotyping approach, usually employed for protists such as *Tetrahymena* spp (Jacob *et al*. 2019; Raffard *et al*. 2023), with a collection-scale analysis focused exclusively on the free-living stage of the pathogen in the absence of host-derived cues to determine whether phenotypic traits are intrinsically organized into an exploration syndrome. Beyond increasing experimental throughput, our approach enables the quantification of behavioural traits across large strain collections, traits that have rarely been incorporated into plant-pathology studies.

Collectively, our findings support the existence of an exploration syndrome underlying the capacity of *A. euteiches* to explore its environment before host contact. This ‘exploration’ syndrome shows that (i) strains with extensive mycelial occupancy developed dense mycelial networks alongside with a high zoospore dispersal rate, and (ii) that strains with higher asexual reproductive output produced smaller and elongated zoospores with faster and straighter swimming trajectories. We identified different combinations of these components among groups of strains in the collection and propose that the three robustly supported clusters represent alternative exploration strategies of the pathogen. The association between phenotypic clusters and the geographic origin of the strains suggests that alternative exploration strategies have emerged during the evolutionary history of *A. euteiches*. The North American cluster (#2) was characterized by a low mobility zoospore exploration strategy, combining rounder and larger zoospores with reduced swimming performance and less linear trajectories. This strategy may reflect adaptation to contrasting edaphic environments since it is known that *Phytophthora* sp. zoospore movement is influenced by soil texture, chemical properties and water availability (Bassani *et al*. 2020). One possible evolutionary scenario would be that in highly structured or poorly drained soils, reduced swimming speed and less linear trajectories may favour the local exploration of bigger zoospores (more reserves) outcompeting foreign zoospores more phenotypically adapted to long-distance exploration. Further analyses on the association between soil structure and the described exploration syndromes are required to validate our hypothesis. In such case, it would open the possibility that differences in soil physical properties, together with contrasting agroecosystem management practices, including perennial forage cropping and crop rotation, may have contributed to the observed phenotypic divergence between North American and European strains.

European clusters (#5, #6) exhibited the greatest phenotypic heterogeneity, particularly for sporulation and mycelial development traits, revealing two contrasting exploration strategies. These strategies rely on different combinations of the two components defining the exploration syndrome. The high-mobility zoospore exploration strategy may promote the rapid exploration of numerous small zoospore swimming fast and linearity. This pattern was found in other oomycetes, such as *Phytophthora parasitica* (Tran *et al*. 2022), suggesting trans-species strategies of zoospore motility. We also observed the common trade-off linking offspring size and number (Smith & Fretwell 1974). High sporulation capacity at the expense of cell size in oomycetes may represent an adaptation to ephemeral, frequently and/or heavily disturbed habitats that necessitate high propagule production to ensure clones’ survival.

In contrast the mycelium-dominated exploration strategy was characterized by extensive mycelial occupancy associated with dense mycelial networks. Although mycelial architecture was not quantified directly in our study, this strategy resembles the phalanx–guerrilla continuum described for fungal mycelia (Aleklett *et al*. 2021). In this framework, phalanx strategies maximize local resource exploitation through dense, highly branched mycelial networks, whereas guerrilla strategies favour exploratory growth and dispersal. Our results suggest that *A. euteiches* similarly exhibits alternative exploration strategies that balance between mycelial and zoospore exploration according probably to environmental conditions and agricultural practices. Intriguingly, high and dense mycelial occupancy was positively associated with high zoospore dispersal rate. Such correlation between two free-living stages of a pathogen pleads for a better integration of its entire lifecycle, and opens the possibility that the willingness to leave a natal habitat as a zoospore is controlled by the same mechanisms as the willingness to explore the soil with a filamentous apparatus. A next step consisting in the molecular investigation of this question seems promising. It is noteworthy that morphological and behavioural attributes were not linked to dispersal rate at the collection scale. This contrasts with a previous experiment describing a cross-species dispersal syndrome linking morphology and activity to dispersal ability (Cote et al. 2022). However, it is also possible that strain-specific phenotypic strategies exist, for instance some strains dispersing as big cells, other as small cells, therefore masking a global relationship. Such strain-specific effects have already been described in ciliates (Junker et al. 2021) and should be investigated oomycetes.

Overall, we describe important phenotypic variation that could reflect genetic adaptation in *A. euteiches*. It is also possible that phenotypic plasticity plays an important role in the expression of the phenotypic strategies we have highlighted here. Distinguishing between genetic and plastic mechanisms in this context is an important research avenue as it will help to inform on the stability of the pathogen exploration structures in the face of different plants and soils.

*In vitro* infectivity assay identified significant positives relationships between mycelial phenotypic traits and pathogenicity while no correlation was associated with *in vivo* infectivity. These positives correlations show that phenotypic mycelial traits participate to explain the *in vitro* infectivity score. Strains with higher occupancy and dense mycelial development show also higher *in vitro* infectivity score under our measurement condition. Moreover, PERMANOVAs indicate relationships between strain pathotype differentiation and to a lesser extend *in vitro* infectivity with exploration syndromes. This study shows for the first time in *A. euteiches* that strain functional traits are associated with pathogenicity and that trait-based approaches could be useful to understand pathogenicity variation in a species.

The lack of correlation between the two infectivity assays further suggests that they reflect distinct components of the infection process. As discussed above, most pathogenicity assays evaluate disease development after the pathogen has already reached or penetrated the host. Our findings extend this view by showing that phenotypic traits expressed before host contact may also contribute to infection success.

This study provides a first functional framework for understanding exploration strategies in *A. euteiches*, nevertheless several limitations should be considered.

First, phenotypic traits were measured under standardized laboratory conditions using a single culture medium and a fixed temperature (22°C). However, prior work have shown that mycelial growth and biomass production vary according to nutrient composition (Papavizas & Davey 1960), while thermal preferences differ among *Aphanomyces* species (Fowles 1976). Consequently, the phenotypic diversity described here likely represents only part of the ecological variation. Future studies should assess the phenotypic plasticity of these traits across a broader range of conditions with a particular emphasis on thermal variation, to better reflect natural environmental variation.

Second, our analyses focused on traits associated with the free-living asexual stage of the pathogen, with zoospore traits quantified as a single endpoint. However, zoosporogenesis is a highly dynamic process. For example, in *Aphanomyces astaci*, zoospores released from hyphae undergo repeated emergence and encystment cycles before locating a suitable host (Francesconi *et al*. 2024). Similar dynamics are likely to occur in *A. euteiches*, although they remain poorly characterized (Beakes *et al*. 2012; Hoch & Mitchell 1972, 1975). Future studies should quantify these temporal dynamics, together with additional functional traits such as flagellar morphology and swimming behaviour (Tran *et al*. 2022). More broadly, our study did not address variation in sexual reproduction, despite evidence that reproductive traits differ substantially among *A. euteiches* strains (Davey & Papavizas 1962, Billard et al., 2018). Oospore production and viability can be quantified both in infected roots and also on synthetic media (Billard *et al*. 2019), making these traits readily accessible for future phenotypic studies. Their integration would also be valuable in light of growing evidence that outcrossing may occur in *A. euteiches*, despite its predominantly homothallic reproductive mode. Adding both asexual and sexual life-history traits would provide a more comprehensive characterization of the performance of *A. euteiches* and further refine the exploration syndrome described here.

Finally, expanding the strain collection to include a broader diversity of geographic origins, hosts and pathotypes will be necessary to better capture the ecological and evolutionary diversity of *A. euteiches*. In particular, phenotyping traits in additional North American pathotype I strains (e.g. MV1, MV5 and MV7) collected from the same nursery as the pathotype III isolate MV3 (Mount Vernon, this study) would provide a valuable opportunity to disentangle the effects of pathotype and environment on the expression of exploration syndrome and infectivity. Beyond ecological diversity, broader sampling will also be essential to resolve the evolutionary history of *A. euteiches*. Kälin et al. (2022, 2026) showed that Italian populations are genetically distinct from other European populations and proposed that they originated from a recent introduction of non-European lineages. Consistent with this hypothesis, a preliminary whole-genome analyses indicate that Italian strains cluster more closely with North American than with other European strains (data not shown). Future studies incorporating the recently identified Canadian strains from pea-growing regions (Gali *et al*. 2026; Sivachandra Kumar *et al*. 2021), together with a broader collection of United-States strains, will be valuable for testing the hypothesis of transcontinental introductions and refining species boundaries within *A. euteiches*.

In conclusion, our study provides evidence for the existence of an exploration syndrome in *A. euteiches*, describing coordinated variation in free-living phenotypic traits before host contact. This syndrome is expressed through alternative exploration strategies that may influence host encounter, the first step of the infection process. Integrating free-living phenotypic traits into studies of soil-borne plant–pathogen interactions could improve our understanding of pathogen adaptation and evolution, while opening new opportunities to develop disease management strategies that reduce pathogen threats by disrupting pathogen exploration before host contact.

## Acknowledgments

We thank Magnus Karlsson and Anne Moussart for sharing strains of *A. euteiches*. We thank Joan Fargues for its laboratory assistance with the *in vitro* infection assays. We thank Aurélie Le Ru for help in imagery analysis. We thank Pablo Molina Vila for its technical assistance in the use of Plateau Technique des Laboratoires – microbiologie, etc. at the SETE, CNRS (UAR 2029). We thank Mohamed Zouaoui for help in whole genome analysis.

Financial support was provided by grants from the “Laboratoires d’Excellences (LABEX)” TULIP (ANR 10 LABX 41), the “Ecole Universitaire de Recherche (EUR)” TULIP GS (ANR 18 EURE 0019) and the Défi-Clé Octaave of the French Région Occitanie.

## Author contributions

MT, DL and EG designed the study and performed experiments. MT performed statistical analysis with contributions from DL and EG. MT, DL and EG wrote the manuscript.

MLPN participated in the collection establishment and supervised the *in vivo* infection assays. CL performed the *in vivo* infection assays. BM performed libraries construction and the nanopore genome sequencing. JG led the genome assemblies. SB built the in-house data-base *Aphanomics* and uploaded the genomes online. HP led the distance-tree analyses.

All authors approved the final version of the manuscript.

## Data availability

Raw data bases and R code for analyses will be accessible online on Zenodo upon the publication of the article.

